# Wind pattern oscillations explain seabird movements at-sea: a nested multiscale approach

**DOI:** 10.64898/2026.04.01.715798

**Authors:** Amedee Roy, Karine Delord, Christophe Barbraud, Pascal Terray

**Affiliations:** Institut de Recherche pour le Développement (IRD), Marbec (Université De Montpellier, IFREMER, CNRS, IRD), Sète, France; Centres d’Etudes Biologiques de Chizé UMR7372, Centre National de la Recherche Scientifique, La Rochelle Université, Villiers en Bois, France; Laboratoire d’Océanographie et du Climat: Expérimentations et Approches Numériques, Institut Pierre-Simon Laplace, Sorbonne Université/CNRS/IRD/MNHN, Paris, France

**Keywords:** south Indian Ocean, climatology, Amsterdam albatross, animal movement

## Abstract

Wind has a strong influence on the flight characteristics, movements, energetics, demography, life-history traits and biogeography of flying animals. With climate change affecting atmospheric circulation patterns at different time scales, understanding the links between wind and animal movements is crucial for predicting its impact on flying biodiversity. Most studies on the relationship between wind and seabird movements have, however, focused on local scales, exploring birds’ perceptive sensitivity to local wind.

In this study, we examine low-level wind pattern oscillations in the Southern Indian Ocean at multiple time scales to explain the local- to large-scale movements of the Amsterdam albatross. Adult individuals exhibited smooth trajectories, strongly correlated with seasonal, intra-seasonal or interannual wind oscillations. Conversely, younger individuals displayed more erratic and exploratory movements, often being swept away by eastward moving low-pressure systems at a synoptic time scale.

Our results suggest that Amsterdam albatrosses can learn and adapt to the annual and monthly low-level wind climatology and interannual variability of the Southern Indian Ocean. This also highlights the importance of investigating seabird movements in relation to broader-scale wind patterns to support their conservation in a changing climate due to human activities. A robust assessment of regional circulation response to climate change for upcoming decades could help project the impact of climate change on seabird movements and mitigate its effects.

## Introduction

Surface winds over lands and oceans have changed during the past decades with marked regional variations across the globe (Young and Ribal 2019; Sharmar et al. 2021; Blunden et al. 2023). Moreover, many of these atmospheric circulation signals have been attributed to anthropogenic forcing (Shaw et al. 2024), including a widening of the Hadley cell and a storm track strengthening in the Southern Hemisphere (Shaw et al. 2022; Lionello et al. 2024; Kang et al. 2024).

Wind has a strong influence on the flight characteristics, movements, energetics, demography, life-history traits and biogeography of flying animals (Drake and Farrow 1988; Chapman et al. 2011; Weimerskirch et al. 2012; Safi et al. 2013; Thorne et al. 2023). As climate change is having an impact on various atmospheric circulation patterns, including the intensity and location of prevailing winds, and that these changes are expected to accelerate in the future (Cai et al. 2005; Sydeman et al. 2014; Masson-Delmotte et al. 2021; Shaw et al. 2022), it is crucial to understand the links between wind direction and intensity and animal movements at process levels, but also at larger spatial and temporal scales to better predict the impacts of climate change on flying biodiversity (La Sorte and Fink 2017; Skyllas et al. 2023).

At fine scale (10s-100s of meters and minutes-hours; Shamoun-Baranes et al. 2017), there is a reasonable understanding of how wind speed and direction influence flight speeds, behaviour and energetic costs of individuals (Pennycuick 1978; Alerstam et al. 1993; Weimerskirch et al. 2000; Amélineau et al. 2014; Van Doren and Horton 2018). However, the influence of wind conditions on flying animals at synoptic and larger spatial and low-frequency time scales has been far less studied and remains poorly understood. Recent studies, based on explicit analytical frameworks, have revealed the influence of broad-scale weather conditions on landbirds stopover distributions (Clipp et al. 2020), high altitude migration (Dokter et al. 2013) and nocturnal migration (Manola et al. 2020; Roy et al. 2025) in terrestrial birds, and migration is a few species of insects (Johnson 1995). Surprisingly, little attention has been paid to the response of seabirds to synoptic scale weather systems (Thorne et al. 2023), despite the long recognized importance of wind for these species, dating back to Leonardo da Vinci (Richardson 2019), and the increasing availability of data on seabird movements (Birdlife International 2023 – Seabird tracking database). Earlier attempts to investigate relationships between seabird movement and synoptic weather patterns based on at-sea observations (Spruzen and Woehler 2002) suffered from mismatches between spatial and temporal scales of weather and seabird data, or had limited time scale and number of individuals monitored (Adams and Flora 2010). Here, to fill this gap, we aim at quantifying the association of seabird movements at sea with winds at various spatial and temporal scales encompassing the regional-, meso- and synoptic scales.

Linking movements at sea to wind patterns is required for understanding aspects of foraging ecology and migration, especially among far-ranging seabirds such as petrels and albatrosses. Previous studies have evidenced the optimal use of wind by seabirds for long-range movements (Weimerskirch et al. 2000; Hromádková et al. 2020; Ventura et al. 2020; Kürten et al. 2025). Wind fields inherently determine the energy expenditure of many seabird movements, impacting their flight behaviour and thereby ultimately leading to changes in population distribution and demographic processes (Spear and Ainley 1998; Péron et al. 2010; Weimerskirch et al. 2012; Ventura et al. 2020). However, to our knowledge, no previous study has attempted to assess the contributions of multiscale wind patterns to seabird movements. Moreover, for birds, movements and behaviours are known to change with age. For example, juveniles birds during the post-fledging period are reported to undertake vagrant erratic journeys and to exhibit changes in behaviour compared to adults (de Grissac et al. 2016a; Brønnvik et al. 2024). Young naïve individuals combine lack of experience and physical immaturity resulting in possible inadequate/inferior skills (Riotte-Lambert and Weimerskirch 2013; Sergio et al. 2022; Delord et al. 2024; Fisel et al. 2024). Yet, whether wind patterns at synoptic or larger scales affect age classes differentially remains poorly known so far.

The Amsterdam albatross *Diomedea amsterdamensis* is a large and long-lived pelagic seabird with an extended immaturity stage (∼ 9 years, (Rivalan et al. 2010)). Like other great albatrosses, it is a biennial breeder (Roux et al. 1983; Jouventin et al. 1989), i.e. adults that raised a chick successfully do not start a new breeding cycle after chick fledging, but remain at sea for a sabbatical period (∼1 yr, (Rivalan et al. 2010)). Its foraging strategy relies on very low flight costs as a result of their dynamic soaring flight, whereby individuals optimize the orientation of their movement with wind direction to maximize the daily distance covered (Pennycuick 1982). During breeding, Amsterdam albatrosses generally head westwards from their colony into the Crozet Basin area until reaching or crossing the Southwest Indian Ridge (Thiebot et al. 2014). During their post-breeding sabbatical period adults move widely (31° to 115° E), mostly exhibiting westwards wider-scale migratory movements (sensu Weimerskirch et al. 2015a) reaching >4000 km from the colony (Thiebot et al. 2014). Immature birds moved widely in longitude (0° to 135° E) and juveniles exhibit very large migratory capacities over the Southern Indian Ocean after fledging (15° to 135° E, ∼ 4500 km from the colony, (Thiebot et al. 2014; Delord et al. 2024).

In this study, we assess how Amsterdam albatrosses respond to multiscale wind patterns across the Southern Indian Ocean over the entire year. Specifically, we investigate (i) the wind climatology and wind pattern oscillations at the regional scale and at multiple time scales, (ii) the relationships between wind patterns and bird movements, (iii) the contributions of wind patterns to albatrosses’ movements, (iv) whether these relationships and contributions vary between age classes. Based on previous studies relating differences in at-sea distribution and behaviour of different life stages (juveniles, immatures and adults; (Thiebot et al. 2014; de Grissac et al. 2016b; Delord et al. 2024)), we expect age classes to differ in their response to wind pattern oscillations. The novelty of this research lies in using a nested multiscale modelling approach of wind patterns in order to understand how these patterns affect seabird movement at sea.

## Materials & Methods

### Tracking data

A total of 36 Amsterdam albatrosses (10 juveniles, 11 immatures and 15 adults) were instrumented in 2006, 2009, 2011 and 2012 with geolocator trackers at the Plateau des Tourbières colony, Amsterdam Island (37° 50’ S; 77° 33’ E) situated in the subtropical part of the Southern Indian Ocean (see supporting information, Tables S1, S2, S3). These sensors recorded time-series of light intensity as well as proportion of time flying. Recorded time-series of light intensity were derived into time-series of longitudinal movements using the dedicated GeoLightR package (Lisovski and Hahn 2012). For more details about animal handling, logger deployment/recovery, and the data processing of geolocator, see the supporting information. It is worth to note that no latitudinal information of albatross movements was used in this study knowing that Amsterdam albatrosses have a relatively low latitudinal range (Thiebot et al. 2014, 2016) and that latitudes estimated from light-based geolocation have lowest accuracy (Phillips et al. 2004; Lisovski et al. 2012).

### Indian Ocean wind patterns

#### Surface wind data

We obtained 6-hourly surface wind data for the 2006-2013 period from the recent state-of-the-art and high-resolution CERSAT ocean surface wind product, which is a blend of satellite observation and the most recent atmospheric reanalyses (E.U. Copernicus Marine Service Information 2012). Zonal and meridional wind components were then extracted for the region from 0°E to 150°E and from 50°S to 20°S and averaged daily to produce daily zonal and meridional wind time-series for assessing the wind-albatross relationships. We further decomposed this daily surface wind variability in three different time scales, which were associated with very different physical mechanisms and processes in the climate system (see below) in order to disentangle their specific and relative roles in albatross movements: the annual (a strictly periodic 1-year signal), low-frequency (>60 days, after excluding the periodic annual signal) and high-frequency (<60 days) time scales.

The wind climatology (i.e., the annual time scale) was obtained, first, by yearly averaging of the daily wind time-series over the 2006-2013 period and, second, by smoothing in time this raw wind climatology. The smoothed wind climatology was estimated through a sequence of applications of locally weighted regression or low-order polynomial (e.g., Loess) to data windows of 80 days applied to the raw wind climatology (Cleveland and Devlin 1988). To estimate the low- and high-frequency wind variability, we then proceeded in two steps. We first removed the annual component by subtracting the smoothed climatology from the original wind time-series and these wind anomaly time-series were finally split into low- and high-frequency components by windowed filtering (Iacobucci and Noullez 2005). The windowed filter was applied to the time-series in the frequency domain. The filter was obtained by convolving a raised-cosine window with the ideal rectangular filter response function. This filter is stationary and symmetric, therefore, it induces no phase-shift and is a good candidate for extracting frequency-defined series components from short-length time-series.

#### Wind pattern oscillation indexes

As our goal is to assess the relationships of wind variations and bird movements at the regional scale, and not at a local scale, we used empirical orthogonal functions (EOF), also known as Principal Component Analysis (PCA), to extract the dominant modes of space-time variability from surface wind patterns at the different considered time scales (i.e. annual, low- and high-frequency). EOF is a popular reduction dimension tool in climate science, which aims at describing the evolution of a variable in space and time (Hannachi et al. 2007; Jolliffe 2011). This technique is notably used to derive oscillation indexes describing the variability of large-scale oceanographic or atmospheric circulation patterns, such as the North-Atlantic oscillation, Southern Annular Mode and the El Niño Southern Oscillation (Rasmusson and Wallace 1983; Jianping and Wang 2003). The key idea of EOFs is to break down a spatio-temporal dataset into few modes of spatial-time variations (i.e. eigenvectors spatial patterns and their associated principal components describing their amplitude and sign variations in time). For a given time, any spatial (wind) map can then be described by few eigenvectors and the amplitude of the associated principal components for this given time. The corresponding eigenvalues define the relative importance of each leading mode of variation for the whole dataset. Here, we applied EOF separately on annual patterns, low-frequency and high-frequency anomalies of zonal and meridional wind components considering two leading modes of variation for each wind component and time scale. This led to the definition of 12 oscillation indexes describing the Indian Ocean surface wind patterns at multiple time scales.

### Nested multiscale modelling of movement patterns

To assess the contributions of wind patterns to albatrosses’ movement at different spatiotemporal scales, we developed a three-step nested modeling approach. We split each time-series of longitudinal movements into several tracks of length 160 days, and for each of these tracks, longitudinal movements were modelled following a hierarchical modeling process with three nested linear models:

- A first linear model was used to explain longitudinal movements using annual pattern oscillation indexes (i.e., the main modes of the EOFs for both zonal and/or meridional wind climatologies derived above)
- A second linear model was used to explain the residuals of the first model using low-frequency oscillation indexes
- A third linear model was used to explain the residuals of the second model using high-frequency oscillation indexes

At each step, we tested multiple linear models with distinct explanatory features. Simple models were considered with distinct combination of the main modes of the EOFs for both zonal and/or meridional wind components. More complex models were considered with possible combination of second-order terms for nonlinear effects as well as cross-terms between variables. All of these models were trained and evaluated through 10-fold cross-validation. It consisted in repeating the following procedure 10 times:

- Splitting the considered track into two distinct training and validation sets of longitudinal position. We considered 70% and 30% of positions for training and validation sets, respectively.
- Fitting the linear models with distinct combination of input features on the training dataset
- Evaluating each linear model using R² score on the validation dataset

The final model for each scale was selected based on the highest average R² score across all validation sets. It was then finally refitted to whole track, and the R² score was used to describe the quality of the fit. This nested approach resulted in three scale-specific linear models for each track.

## Results

### Indian Ocean wind patterns

#### Surface wind annual patterns

At the annual time scale, the surface wind variations over the region were dominated by the Mascarene High (MH), which is a semi-permanent subtropical anticyclone (Krishnamurti and Bhalme 1976; Zhao et al. 2023; Figure 1a) and the Southern Hemisphere Westerlies at higher latitudes. Winds turned anti-clockwise around the MH due to geostrophic balance. Consequently, the surface wind was essentially zonal and westerly below 35°S, but south easterly above 30°S year around (Figure 1a). At the breeding colony latitude (37°50’S), winds were dominantly zonal and eastward and the wind speed increased significantly to the south of the colony, in the westerlies wind regime. The leading EOF modes of the climatological zonal and meridional wind variations with respect to this annual average, which account for about 80% of the variance of climatological zonal and meridional winds, mainly reflected the latitudinal and longitudinal translations of the position and strength of the MH system, generating seasonal wind variations, which repeat each year. As an illustration, at the breeding colony latitude, westerlies were strengthened from May to October (e.g., positive amplitude of the leading eastward wind climatology principal component), but weakened or even became easterlies (e.g., negative amplitude of the leading eastward wind climatology principal component) from November to April (Figure 1b left). At the breeding colony latitude, the meridional climatological wind variations were small year around as the spatial loading at this location in the meridional climatological wind was almost zero (Figure 1b right). The seasonal meridional wind variability was mainly concentrated off Australia and reflects the longitudinal shifts of the MH during the year, with a more westward (eastward) position during austral winter (summer) (Figure 1b, right).

**Figure 1:**
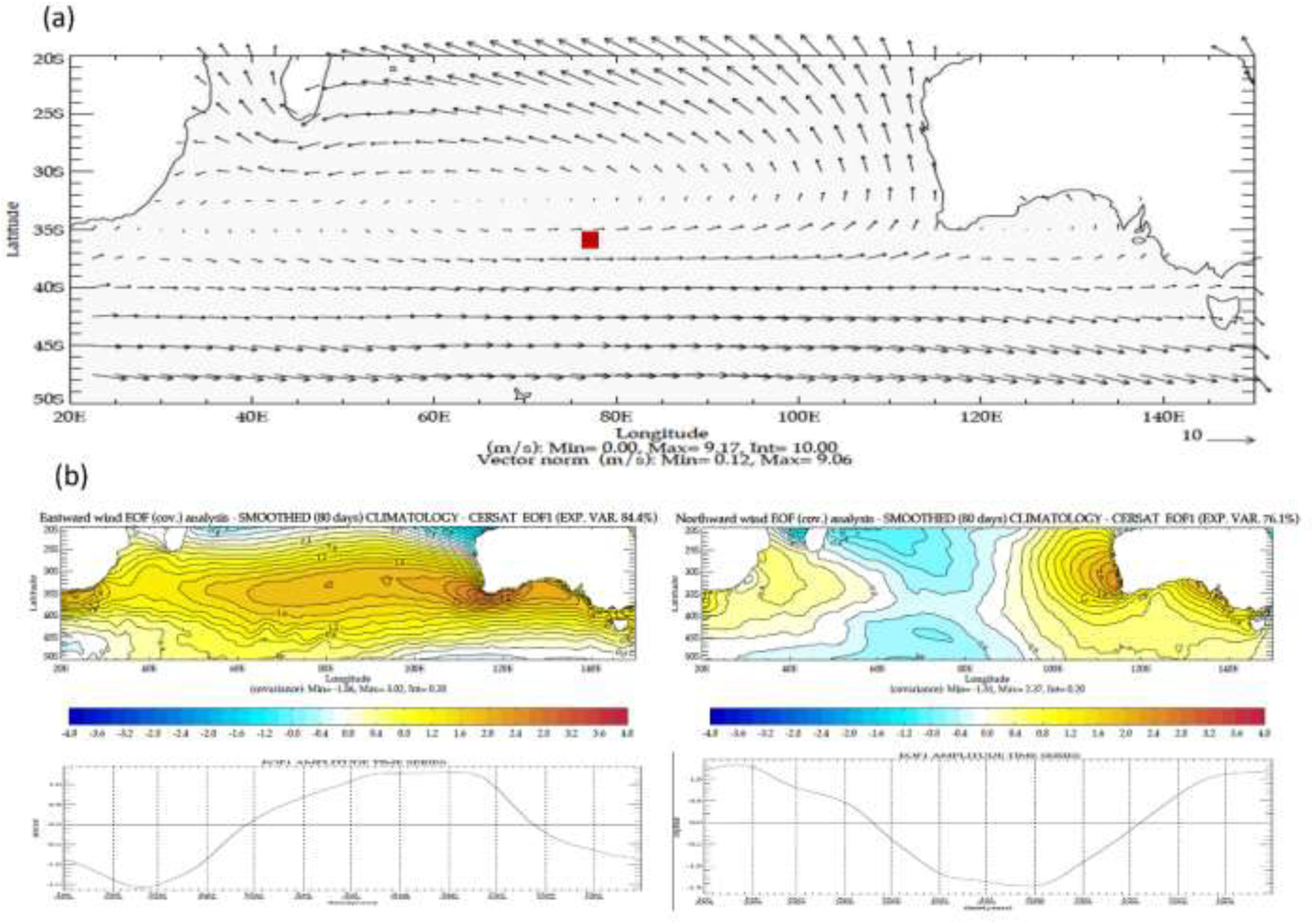
(a) Multi-year average of surface wind over the studied area. The breeding colony of Amsterdam albatrosses is represented by the red square. (b) Main modes of space-time variation of the climatology patterns of eastward (left) and northward (right) winds. Maps describe the main spatial mode derived from the EOF analysis. Time-series illustrate the contribution of these main modes overtime.

#### High-frequency surface wind patterns

The high-frequency surface zonal and meridional wind variability described by their respective two leading modes were typical of an eastward propagating wind system (Wheeler and Hendon 2004), e.g. the two leading modes for each wind component explained roughly the same amount of variance and their spatial patterns were in quadrature of each other (Figure 2a).The associated principal component time-series were uncorrelated simultaneously, but lead-lag correlated (2b). Maximum correlations were obtained for lags of 1 and 2 days for meridional and zonal winds, respectively. In a nutshell, the leading two EOFs of high-frequency wind variations (<60 days) describe the wind variations associated with a succession of the low-pressure disturbances and depressions marking the storm tracks, embedded in the mid-latitude westerlies of the Southern Hemisphere, and travelling across the study area in about 4 to 8 days year around.

**Figure 2:**
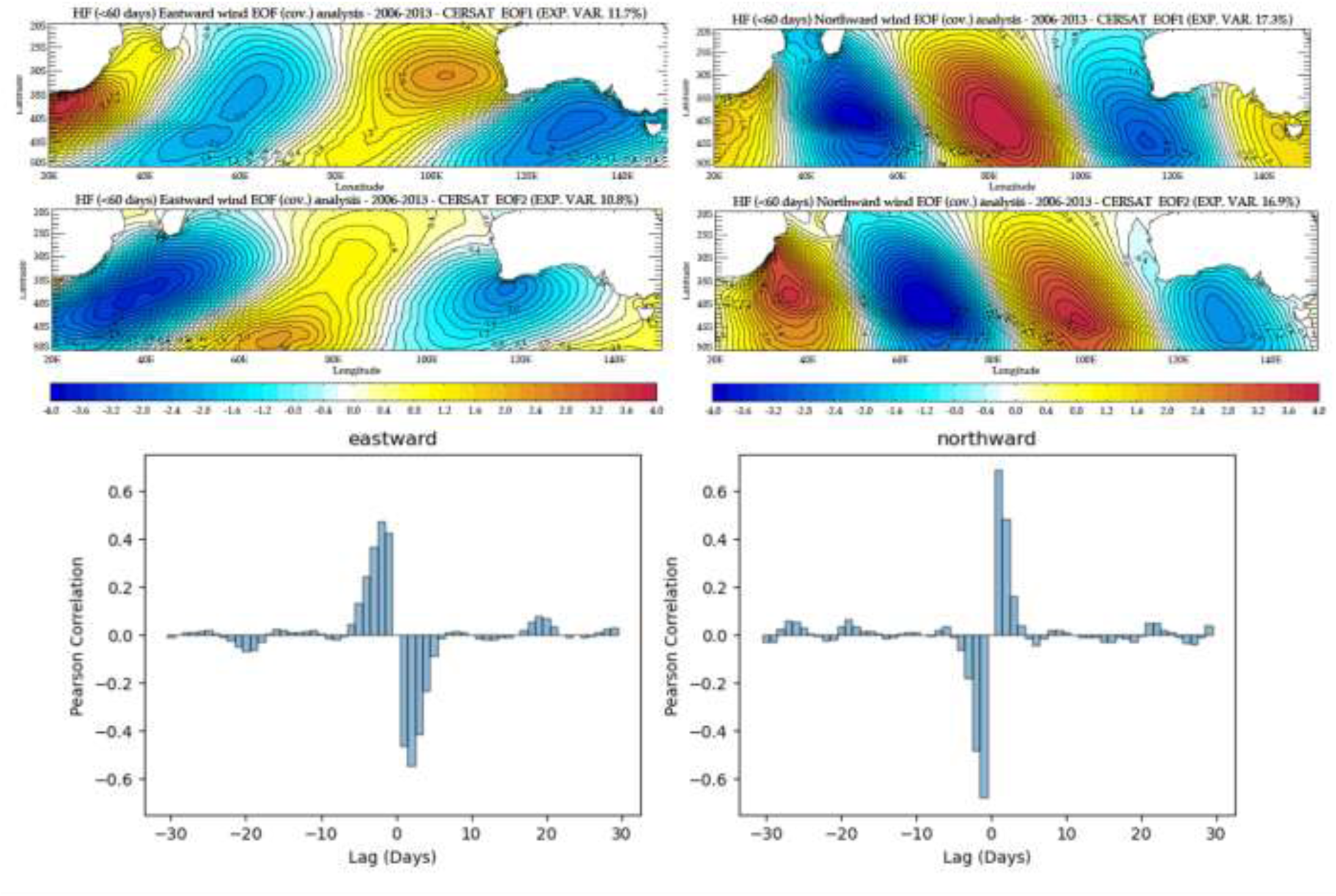
The two leading modes of spatial variation of the high-frequency (< 60 days) patterns of eastward (left) and northward (right) wind components. Maps describe the first two spatial modes derived from the EOF analyses of eastward and northward winds. Histograms illustrate the lagged correlations between the time-series of amplitude variations of the two associated modes for the eastward (left) and northward (right) winds.

#### Surface wind low-frequency patterns

The first two leading modes of the low-frequency zonal wind anomalies were strikingly different from the high-frequency leading modes and were not associated with any obvious space-time propagation of some wind systems (Figure 3). This suggests that these modes represented a modulation of the seasonal cycle at the intra-seasonal time scale, which was well observed on the associated principal components, and, also, perturbations of the position and strength of the MH associated with interannual climate modes, such as the Subtropical Indian Ocean Dipole (SIOD), El-Niño Southern Oscillation (ENSO) or Southern Annular Mode (SAM) phenomena. As an illustration, the second mode seemed to represent north-westward or south-eastward shifts of the position of the MH, which are known to be associated with both ENSO and SIOD (Behera and Yamagata 2001; Terray 2011). In a nutshell, the low-frequency component (>60 days) describes mainly the weather regime shifts associated with interannual phenomena like ENSO or the SIOD, and also intra-seasonal wind oscillations and disturbances.

**Figure 3:**
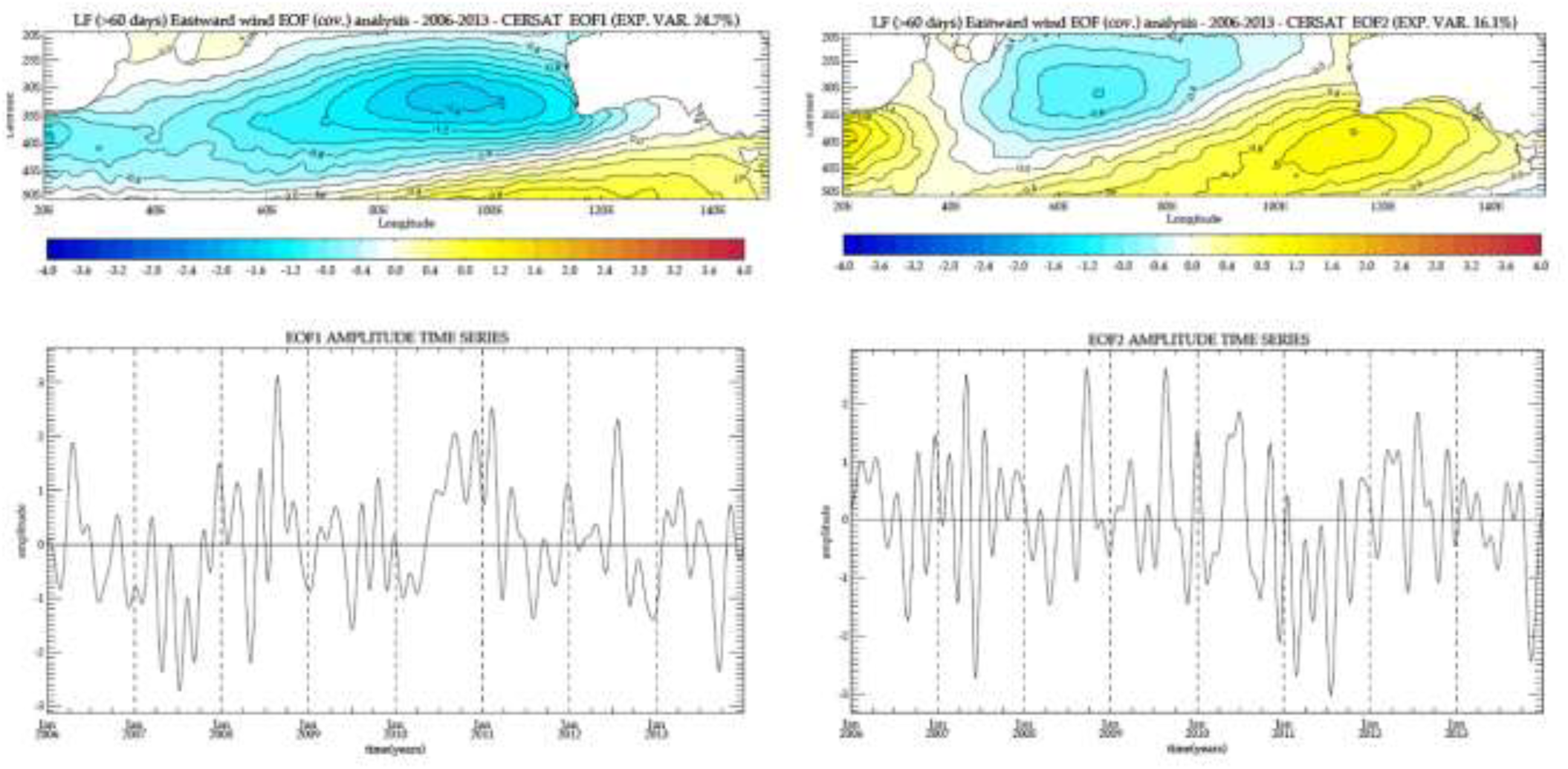
Leading modes of space-time variations of the low-frequency (> 60 days) patterns of eastward (left) and northward (right) wind. Maps describe the main spatial modes derived from the EOF analysis of the eastward and northward wind components. Time-series illustrate the amplitude variations of these main modes overtime.

**Figure 4:**
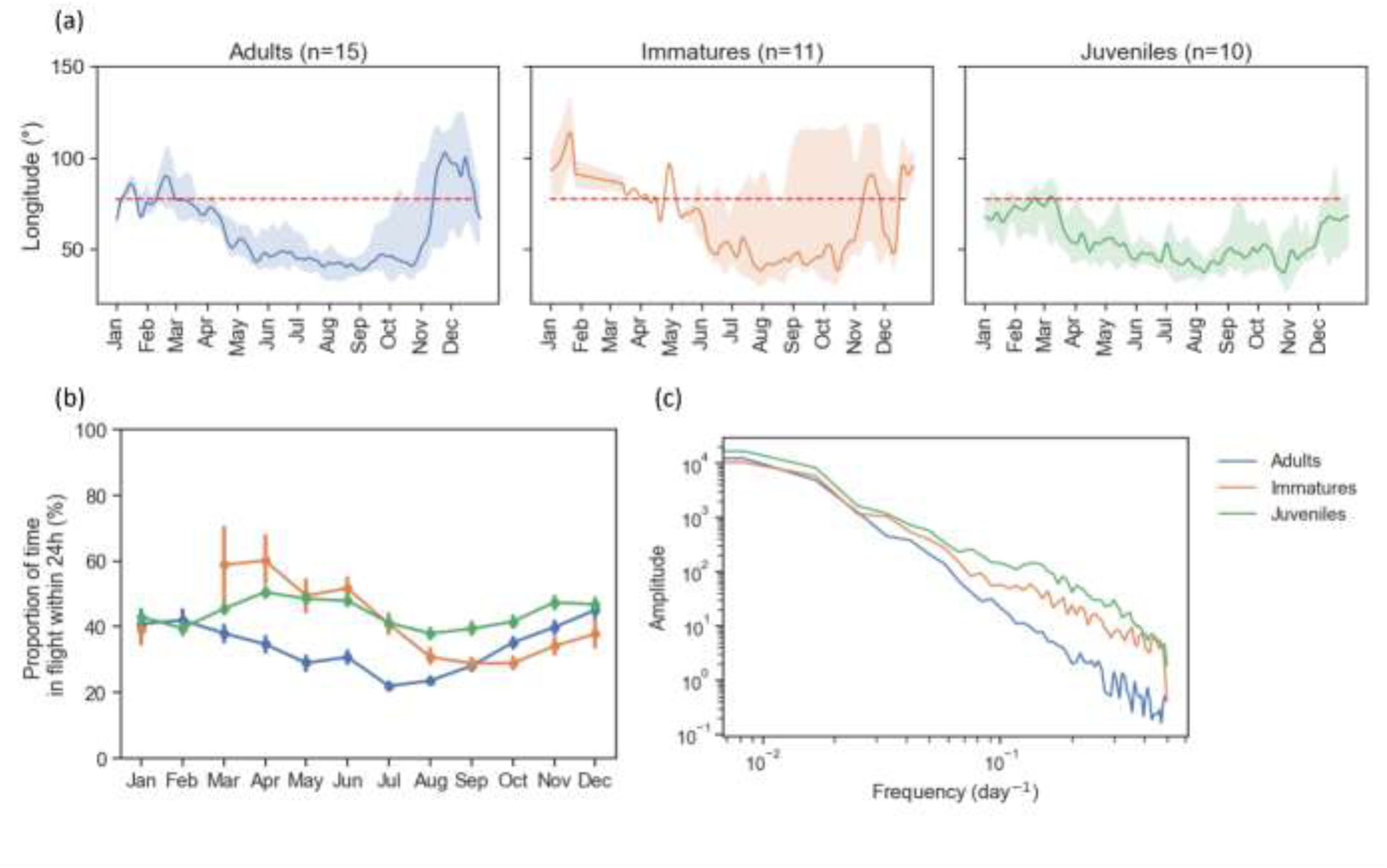
(a) Median longitudinal movements of the Amsterdam Albatross on an annual basis, categorized by stage. Envelopes show 25% and 75% quantiles. The position of the breeding colony is indicated by the red-dotted lines. (b) Average proportion of time spent flying, calculated over 24 hours for each month and stage. (c) Average power spectral density of the longitude time-series, grouped by stage. The color codes are as follows: blue for adults, orange for immatures, and green for juveniles.

### Albatross movements

A total of 36 tracks from the Amsterdam albatross were recorded in 2006, 2009, 2011 and 2012. Trackers recorded trips from 212 to 1220 days, that were split into a total of 75 samples of 160 days (26 adults, 12 immatures and 37 juveniles).

#### General movement patterns

Departure times from the colony differed for adults, immatures and juveniles with individuals leaving, respectively, in early-April, late-May and mid-March. Outside the breeding period, movements ranged from South-African coasts (40°E) to Australian coasts (110°E). Juveniles tended to stay at the west of Amsterdam Island, while adults and immatures oscillated in between west and east of the island. Adults however switched more synchronously than immatures from the western part of the Southern Indian Ocean to the eastern part (Figure a). These variations were also illustrated in terms of proportion of time spent flying (Figure b). Power Spectrum analysis of the longitude time-series demonstrate that they had higher energy at the low-frequency (e.g., seasonal) time scale, reflecting the seasonal patterns in the birds’ movements (Figure c). Interestingly, these spectral densities differed between stages for highest frequency, with highest energy for immatures and juveniles. In other words, younger individuals exhibited more frequent changes in their positions.

#### Stage-specific nested multiscale modelling of movement patterns

The selected linear models incorporating annual oscillation indices explained, on average, 85% of the variance in longitudinal movements of albatrosses. This explanatory power was even higher for adults, reaching 94%. Among the predictors, climatological eastward and northward wind patterns were retained in 98% and 100% of the models, respectively, despite the time series deal only with longitudinal information. Additionally, second-order terms and interaction terms were included in 84% and 98% of the models, indicating a strong non-linear and synergistic influence of wind features at this temporal scale.

Models based on low-frequency oscillation indices explained a more modest 13% of the variance in the residuals from the annual models. For adult individuals, this increased to 22%, although the explanatory power varied widely across tracks—from 0% to over 80%. Low-frequency eastward and northward wind components were selected in 71% and 60% of models, respectively. The inclusion of second-order and interaction terms was less frequent, at 41% and 31%.

Finally, models incorporating high-frequency oscillation indices explained 5.6% of the remaining residual variance. Interestingly, juveniles showed the highest sensitivity to these patterns, with an average R² of 6.9%, and some tracks reaching up to 20%. High-frequency eastward and northward wind features were selected in 66% and 52% of models, respectively. The use of second-order and interaction terms was limited, appearing in only 29% and 4% of models.

The distribution of R² scores by life stage is presented in Figure 5, while Figure 6 illustrates individual-level relationships between wind oscillation patterns and movement trajectories across temporal scales.

**Figure 5.**
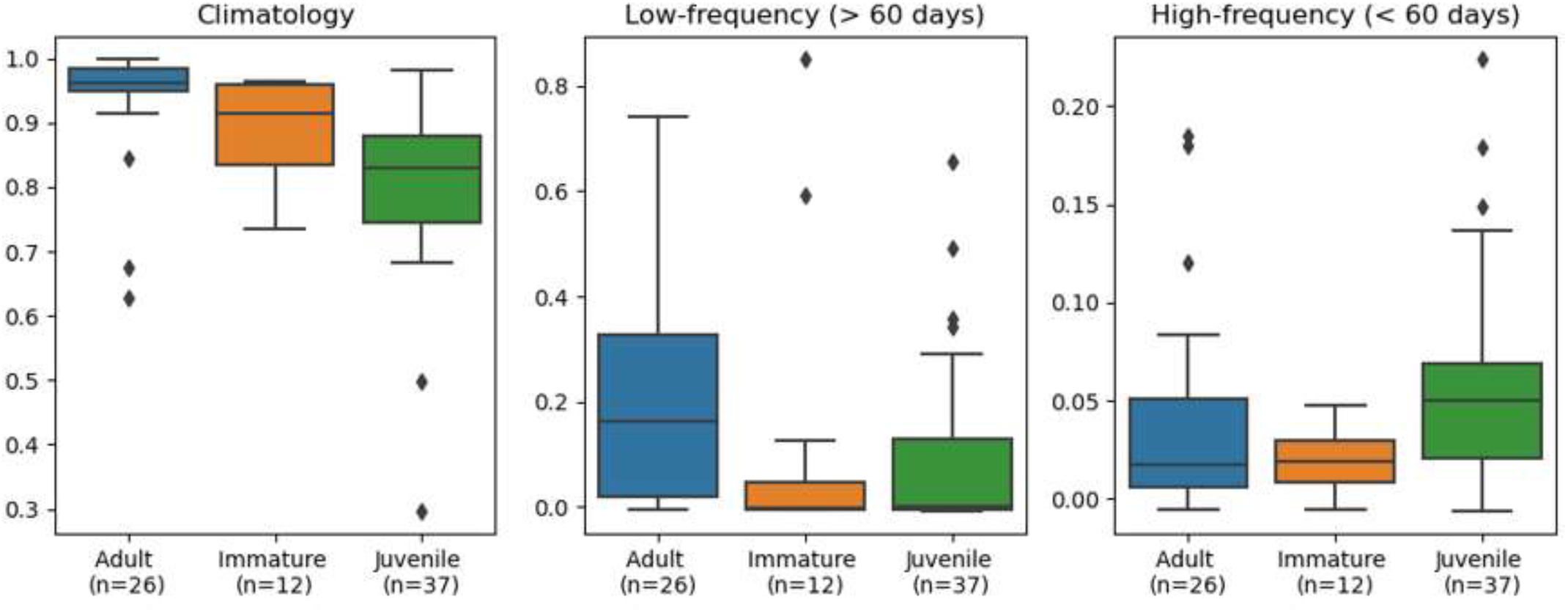
: R^2^ scores of the nested linear models used to explain individual bird movements. Left, middle and right panels describe the R^2^ scores of the linear models dedicated to, respectively, the annual, the low-frequency and the high frequency patterns. R^2^ scores of linear models are grouped into boxplots based on the individuals’ life stages.

**Figure 6.**
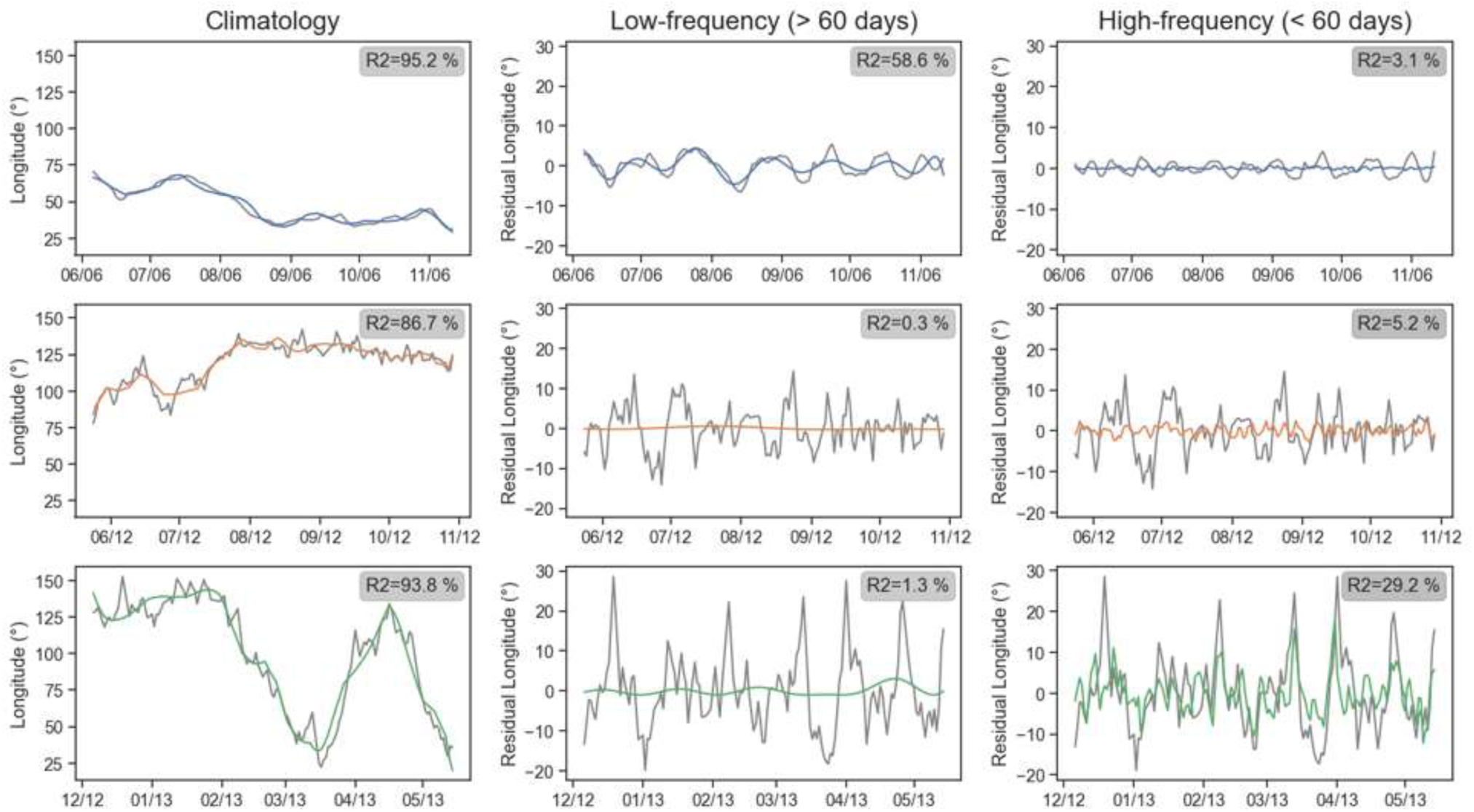
: Example of observed and modelled longitudinal movements for three individuals: BS12376 (Adult/Blue); BS29502 (Immature/Orange); BS29556 (Juvenile/Green). Each column refers to a specific time scale. The first column refers to raw movements’ observation (in grey) and predicted movements from climatological indexes (in color). The second column refers to climatological residuals (in grey) and predicted residual movements from low-frequency indexes (in color). Similarly, the last column refers to residuals of the first two models (in grey) with predicted residual movements from high-frequency indexes (in color).

## Discussion

### Multi-scale effects of surface wind on Amsterdam albatross

This study describes oscillations in wind patterns over the Southern Indian Ocean at multiple spatio-temporal scales and reveals strong relationships between wind oscillations and the longitudinal movement patterns of Amsterdam albatrosses. Our approach decomposes daily surface wind variability into three distinct interpretable components related to different temporal and physical scales: annual, low-frequency (>60 days) and high-frequency (<60 days) time scales. These components relate to various physical mechanisms and processes. On one hand the annual component describes seasonal patterns. On the other hand, the low-frequency component describes mainly the weather regime shifts associated with interannual phenomena like ENSO or the SIOD, and also intra-seasonal wind oscillations and disturbances. Finally, the time variations of the high-frequency component are driven by the succession of the low-pressure disturbances and depressions marking the mid-latitude storm tracks in the Southern Hemisphere.

Interestingly, the movement patterns of the Amsterdam albatross are influenced by wind oscillations across these three different time scales even though the seasonal time scale is the most influential. These movements exhibit a dominant yearly periodicity, as the albatrosses return to their colony at the same time each year. Consequently, their movements are strongly correlated with the annual wind components, with climatological patterns accounting for over 90% of the variance in their longitudinal movements. This synchronicity is particularly pronounced in breeding adults, whose movements are directly tied to their breeding cycle. Adult albatrosses also display reduced erratic movements and a closer relationship to low-frequency wind patterns compared to immature and juvenile birds. This suggests that adults may have developed more efficient flight trajectories over time, reflecting their greater experience of the local climate and its variability at different time scales. They might have learned to deal with regime shifts associated with interannual phenomena or intra-seasonal wind oscillations and disturbances. In contrast, immature and juvenile albatrosses exhibit movement patterns more directly related to high-frequency wind oscillations, reflecting more active or exploratory behaviours. This could be partly attributed to environmental factors, such as wind forcing, with younger birds presumably being more influenced by local weather conditions that cause rapid shifts in their movement patterns. Some high-frequency movement patterns in juveniles were indeed highly correlated with high-frequency wind oscillations, suggesting they are swept by low-pressure disturbances, as illustrated in Figure 6.

### Further understanding of seabird-wind relationship

Amsterdam albatrosses, like other Procellariiform seabirds, adopt a flight behaviour called dynamic soaring, i.e. exploitation of vertical gradients in wind speed near the sea surface at lowest cost (Clay et al. 2020). This limits the directions in which dynamic soaring birds can fly effectively (Ventura et al. 2020). Therefore, there is a need to adjust the search strategies by making trade-offs between knowledge of the location and value of resources and the wind field, which varies in space and time (Thorne et al. 2016). Birds have to cope with both yearly variations and local weather conditions, and resource availability, implying flexibility in movements. Most studies examining the relationship between wind and seabird movements have focused on local scales (Vansteelant et al. 2021). These approaches aim to explain how birds behave locally in the conditions they experience. However, our research indicates that large- to medium-scale movements can also be explained by well-defined (and sometime periodic) oscillations in wind regimes. Exploring the relationships between wind climatology, weather conditions and bird movements at these larger scales reveals other information, which are probably not due to the birds’ perceptive sensitivity, but to their understanding of the dynamics of the climate systems. Low-frequency wind regimes are also likely linked to specific prey distribution and availability, which could explain this synchronicity (Arrigo et al. 2008). This study shows indeed substantial differences in Amsterdam albatross movement patterns related to individual age. This supports the hypothesis that these birds can learn and adapt to the annual and monthly dynamics of the Southern Indian Ocean climate system (Riotte-Lambert and Weimerskirch 2013; Powers et al. 2022).

However, our study does not precisely describe the physical and/or biological processes that the birds may have learned. The tracking data used in this study was limited to longitudinal data due to the inherent latitudinal accuracy of GLS (Phillips et al. 2004; Lisovski et al. 2012). Yet our results suggest that variability of meridional winds have already some significant relationships with longitudinal movements of the birds. More precise information on their latitudinal movements could help to identify the exact areas that adults visited and may lead to a better understanding of the role of the wind forcing at the larger scales. Additional years of data could also help in better identifying interannual weather patterns. Nonetheless, this study demonstrates the benefits of exploring multiscale wind data analysis combined with dimension reduction tools such as Empirical Orthogonal Functions (EOF).

### Impact of climate change on wind patterns & seabird movements

This work matches with three of the recent research priorities highlighted by Thorne et al. (2023) for advancing the understanding of the effects of wind on seabird ecology and behaviour: a) how do seabirds sense and anticipate wind conditions (i.e. ability to use local environmental cues to anticipate wind conditions in space and time), b) what is the role of individual learning on movement responses to wind (i.e. plasticity in behavioural norms and variability among individuals), and c) how understanding of broader-scale wind patterns improve seabird conservation. Albatrosses are of broad conservation concern (Dias et al. 2019) and predictive models of future distribution at sea of these highly mobile species given upcoming climate change often failed to explicitly take into account for wind parameters (Somveille et al. 2020). Our work suggests therefore that exploring seabird movement and wind patterns at a synoptic, interannual and seasonal scales might help forecasting future seabird movements and the impact of climate change.

Seasonal variations in prevailing winds and climatic conditions are somewhat predictable across the globe. The Southern Hemisphere zonal wind stress maximum has increased significantly over the past 30 years and this change can be attributed partly to human activities (Swart and Fyfe 2012; Shaw et al. 2024). Changes in the position and strength of the Southern Hemisphere surface westerlies and the MH have significant implications for ocean circulation and the global carbon cycle (Gent 2016; Goyal et al. 2021). In the Southern Indian Ocean future changes in zonal winds (i.e. the westerlies) are likely with probable consequences on flight efficiency and migration of seabirds (Cai et al. 2009; Goyal et al. 2021; Morten et al. 2023). Species such as albatrosses, that mainly move around the Southern Ocean using soaring flight (Sachs et al. 2013; Schoombie et al. 2023), may have to modify their flight paths to face these changes.

## Acknowledgements

This study was made possible thanks to all the fieldworkers involved in the monitoring program on Amsterdam albatross, namely Jean-Baptiste Thiebot, Jérémy Demay, Rémi Bigonneau, Romain Bazire, Hélène Le Berre, Marine Quintin, Marine Devaud, Chloé Tanton, Jérémy Dechartre and Anthony Le Nozahic. We are grateful to Richard Phillips, British Antarctic Survey, Cambridge for providing GLS loggers. This study is a contribution to the National Plan of Actions for Amsterdam albatross. This monitoring program was supported financially and logistically by the French Polar Institute IPEV (project 109 Ornithoeco, PI C. Barbraud/H. Weimerskirch), the Zone Atelier Antarctique (ZATA, CNRS-INEE), Terres Australes et Antarctiques Françaises. This study is part of the long-term Studies in Ecology and Evolution (SEE-Life) program of the CNRS. All work was carried out in accordance with the IPEV ethics committee. We acknowledge Dominique Joubert for the management of the demographic CEBC Seabirds database. Pascal Terray is funded by Institut de Recherche pour le Développement (IRD, France).

## Data availability

The data and scripts used in this study are available via the InDoRES online data repository: https://data.indores.fr:443/privateurl.xhtml?token=efb20537-3b1b-4485-8327-c0a80da1ec35.

## Statements & Declarations

☒ The authors declare the following financial interests/personal relationships which may be considered as potential competing interests:

Christophe Barbraud reports financial support was provided by French Polar Institute Paul Emile Victor. If there are other authors, they declare that they have no known competing financial interests or personal relationships that could have appeared to influence the work reported in this paper.

## Author Contributions

All authors contributed to the study conception and design. Material preparation, data collection and analysis were performed by Amédée Roy, Pascal Terray and Karine Delord. The first draft of the manuscript was written by Amédée Roy and all authors commented on previous versions of the manuscript. All authors read and approved the final manuscript.

## Supplementary

### Study species, study colony and logger deployments

The Amsterdam albatross, like other great albatrosses, is a biennial breeder (Roux et al. 1983; Jouventin et al. 1989), with high survival during juvenile, immature and adult phase (Rivalan et al. 2010). The adults that raised a chick successfully do not start a new breeding cycle after chick fledging, but remain at sea for a sabbatical period (∼1 yr; Table 1; Rivalan et al. 2010). However, early failed breeders leaved the colony earlier in the breeding season compared to successful breeders, and may start to breed the following year (Rivalan et al. 2010). Immature birds may visit the colony when they are 4−7 yrs old, but generally only start breeding at 9 yrs old (Weimerskirch et al. 1997a). Juvenile birds fledge and migrate independently from the adults in January (Delord et al. 2024). Amsterdam albatrosses were monitored annually since 1983 and all individuals were individually marked (numbered stainless steel and plastic engraved colour bands; see Rivalan et al. (2010) for details).

Amsterdam Island (37° 50’ S; 77° 33’ E) is located in the subtropical part of the southern Indian Ocean. In this oceanic area, the southern subtropical front delimits the warmer subtropical from the colder sub-Antarctic waters (Belkin & Gordon 1996). Though the diet and foraging strategy of Amsterdam albatross remains poorly known, it is presumed to have very similar foraging behaviour compared to that of the wandering albatross, although subtle differences can appear (Pajot et al. 2021). Like other large albatross species (*Diomedea spp.*), the Amsterdam albatross is expected to prey on large squid, fish and carrion found on the sea surface (Delord et al. 2013, Cherel et al. unpublished data). The wandering albatross is known to forage over extensive distances, detecting prey visually or by olfaction during the day (Nevitt et al. 2008).

Thiebot et al. (2014) showed that adult Amsterdam albatrosses during their post-breeding sabbatical period moved widely (31° to 115° E), mostly exhibiting westwards wider-scale migratory movements (*sensu* Weimerskirch et al. 2015a) reaching >4000 km from the colony exploiting continuously warm waters (∼18°C). The immature birds moved widely in longitude (0° to 135° E), exploiting exclusively warm waters 17°-18° C. Similarly to adults no clear longitudinal seasonality synchronicity in the movements was evidenced, except that they also tended to move westwards in June and eastwards in November. Juveniles exhibited very large migratory capacities over the southern Indian Ocean after fledging (15° to 135° E, ∼ 4500 km from the colony), through a large range of latitudinal gradient (27° to 47° S). De Grissac et al. (2016) compared trajectories (i.e. departure direction or orientation toward specific areas) of juveniles and adults and showed that juveniles performed an initial rapid movement taking all individuals away from the vicinity of their native colony, and secondly performed large-scale movements similar to those of adults during the sabbatical period. De Grissac et al. (2016) concluded in an overlap in distribution between adults and juveniles due to the extensive area they used and their differences in latitudinal distribution compared to other Procellariiformes species.

Geolocators-GLS are archival light-recording loggers used to study activity of birds over periods lasting up to ∼ 2 years. GLSs record the ambient light level every 10 min, from which local sunrise and sunset hours can be inferred to estimate location every 12 h (Wilson et al. 1992). GLS loggers allowed us to track the birds for prolonged periods with minimal disturbance to them. We considered the following stages regarding the year of GLS deployment: juvenile, as a fledgling equipped with a GLS just before leaving the colony for the first time; immature, as a non-breeding young bird that had never bred equipped with a GLS when visiting the colony; adult, as a breeding adult equipped with a GLS during the incubation or brooding period which successfully fledged a chick and thereafter took a sabbatical year (data corresponding to breeding period was excluded from the analysis). To date, we have retrieved 36 of the 60 GLS loggers deployed in total over 4 years, from which 36 individual tracks were estimated. Our original aim was to collect data over the three life-stages on a long period of time (>1 year). These data are available from a total of 15 adults tracked throughout their sabbatical period, 11 immature birds and 10 juvenile birds (up to 3.2 years).

### Geolocator data processing

Global Location Sensing (GLS) loggers (British Antarctic Survey, Cambridge) were deployed on individuals during the breeding season (2006, 2009, 2011 and 2012; Tables S1, S2). GLS loggers (see Tables S3) weighted < 0.1% of body mass and well below the recommended threshold of 3% of body mass (Phillips et al. 2003). These were mounted on plastic leg bands. Individuals were captured on the nest during the breeding season.

GLS provided light-level data used to calculate geographic positions. Departure from the colony, hereafter referred as migration, was determined by visual examination of movement; outward migration started from the first directional movement (followed by several consecutive days with directional flight), while the final nonbreeding location was the last location in the non-breeding area before a sustained period of directional movement towards the breeding colony. GLS data allows long-term (several years) latitude and longitude estimation from daylight measurements, albeit although with a lower accuracy (186 ± 114 km; Phillips et al. 2004) than satellite transmitters (Wilson et al. 1992). Loggers measured daylight level intensity every 60 s and the maximum intensity for each 10 min is recorded. An internal clock allows estimation of the latitude based on day length and longitude based on the timing of local midday with respect to Coordinated Universal Time (Afanasyev 2004). Daylight data recorded by GLS were analysed using a standardized procedure for flying seabirds (Phillips et al. (2004)) to provide two locations per day. Recorded timeseries of light intensity were derived into timeseries of longitudinal movements using the dedicated GeoLightR package (Lisovski and Hahn 2012). During a 1-2 wk period around the equinoxes (20-21 March and 22-23 September), latitude cannot be estimated accurately (Wilson et al. 1992).

To select the data corresponding to periods spent at sea after leaving the breeding site, we used the following criteria on activity to define the departure time from the colony for each stage: 1) juveniles, the first bout spent on seawater (wet bouts duration) > 1h based on Argos Platform Transmitters Terminals, PTT tracking data (data obtained in a other project and not shown here, please see Weimerskirch et al. unpublished data); 2) immatures and adults, the last bout spent flying (dry bouts duration) > 12h based on PTT tracking data (Weimerskirch et al. unpublished data). Using these criteria we obtained departure months as follows: 1) the juveniles fledged from the colony in March, 2) the immatures left between May and June, and 3) the departures of sabbatical adults take place between March and April (Table S1).

---

**Table S1.** Summary information for the 36 Amsterdam albatross equipped with geolocator in 2006, 2009, 2011 and 2012, at Amsterdam Island, including year, number of individuals and months of departure.

| Year | Total individuals | Stage | Female | Male | Months of departure |
| --- | --- | --- | --- | --- | --- |
| 2006 | 8 | adult | 4 | 4 | March - April |
| 2009 | 7 | adult | 3 | 7 | March - April |
| 2011 | 7 | immature | 4 | 6 | May - June |
| 2012 | 10 | juvenile | 6 | 4 | March |
| 2012 | 4 | immature |  | 4 | May - June |

**Table S2.**
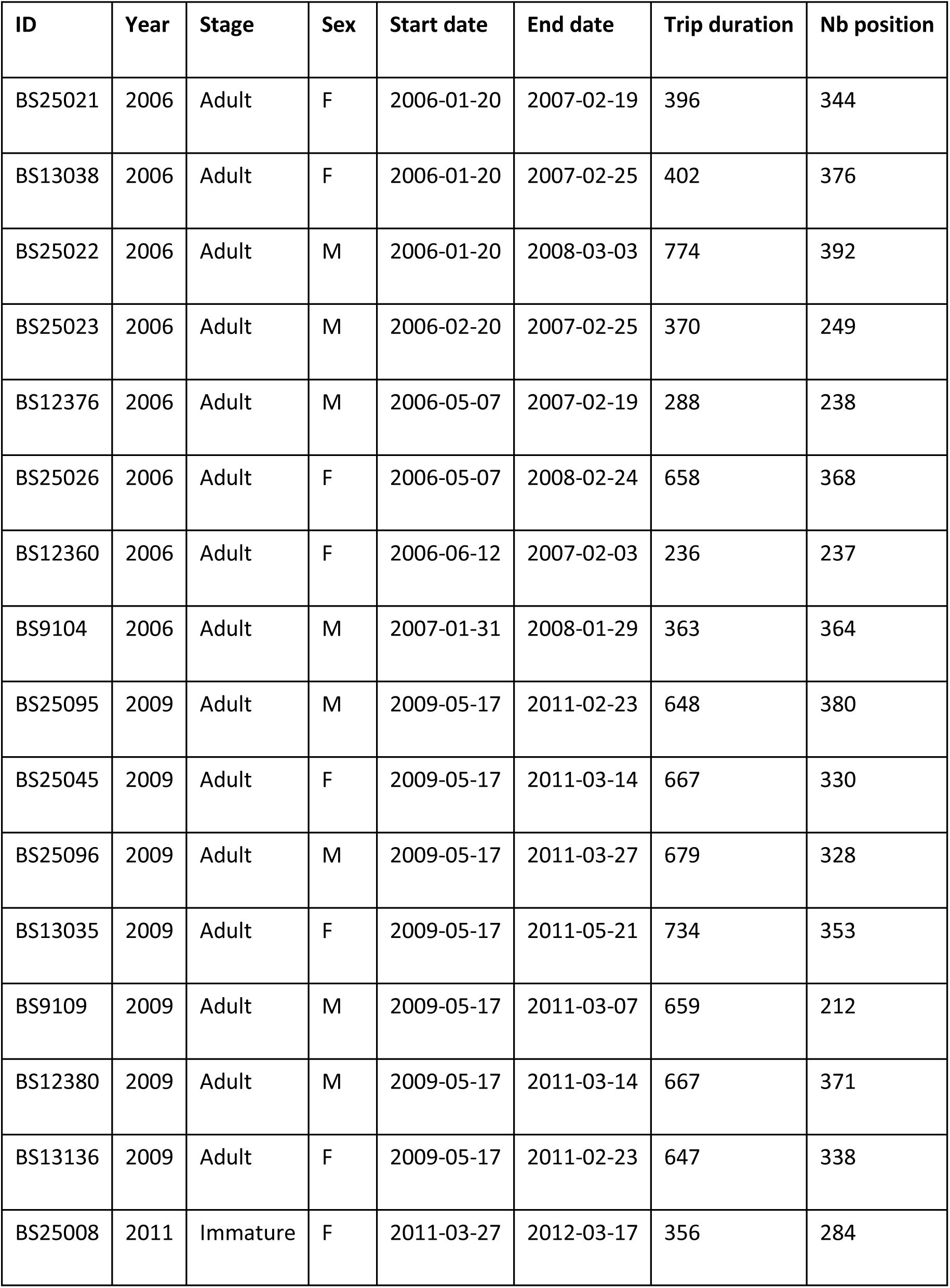
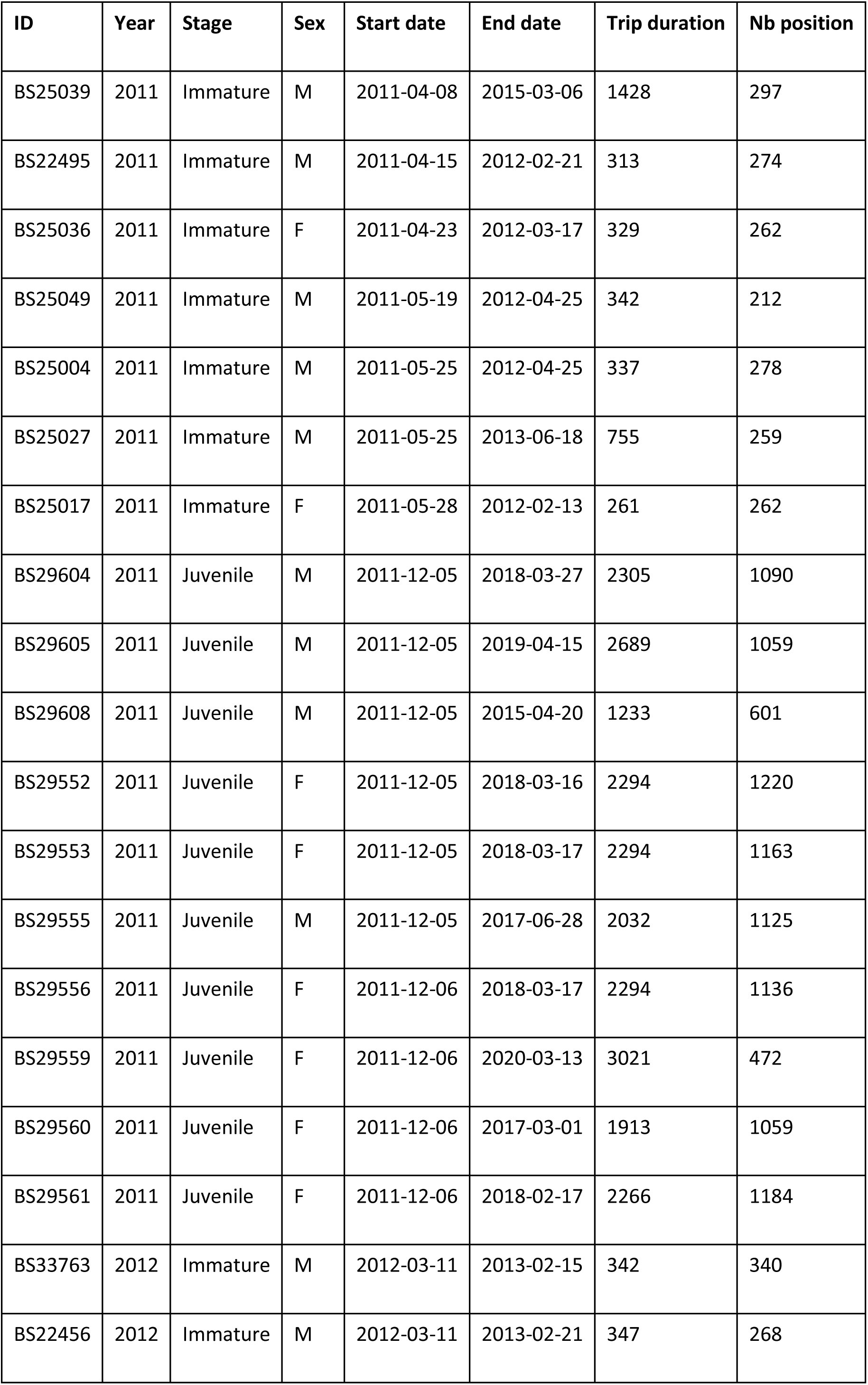
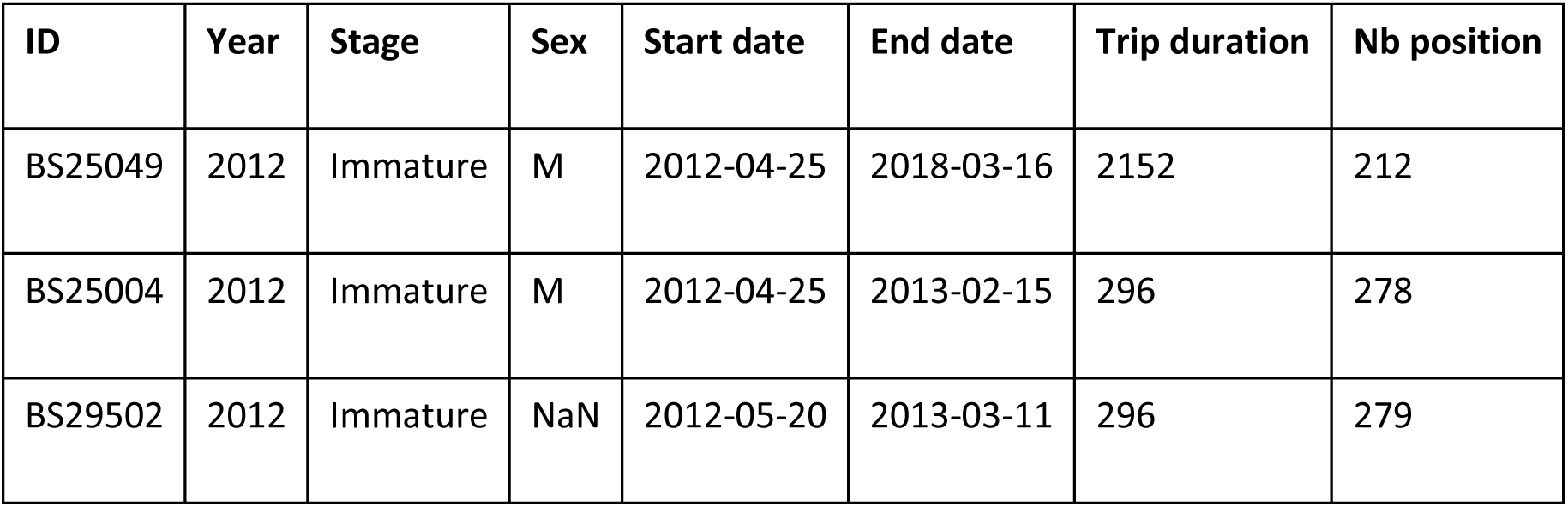
General information on the 36 Amsterdam albatross equipped with geolocator in 2006, 2009, 2011 and 2012, at Amsterdam Island, including year, number of individuals, stage, sex, starting and ending tracking date, trip duration.

| ID | Year | Stage | Sex | Start date | End date | Trip duration | Nb position |
| --- | --- | --- | --- | --- | --- | --- | --- |
| BS25021 | 2006 | Adult | F | 2006-01-20 | 2007-02-19 | 396 | 344 |
| BS13038 | 2006 | Adult | F | 2006-01-20 | 2007-02-25 | 402 | 376 |
| BS25022 | 2006 | Adult | M | 2006-01-20 | 2008-03-03 | 774 | 392 |
| BS25023 | 2006 | Adult | M | 2006-02-20 | 2007-02-25 | 370 | 249 |
| BS12376 | 2006 | Adult | M | 2006-05-07 | 2007-02-19 | 288 | 238 |
| BS25026 | 2006 | Adult | F | 2006-05-07 | 2008-02-24 | 658 | 368 |
| BS12360 | 2006 | Adult | F | 2006-06-12 | 2007-02-03 | 236 | 237 |
| BS9104 | 2006 | Adult | M | 2007-01-31 | 2008-01-29 | 363 | 364 |
| BS25095 | 2009 | Adult | M | 2009-05-17 | 2011-02-23 | 648 | 380 |
| BS25045 | 2009 | Adult | F | 2009-05-17 | 2011-03-14 | 667 | 330 |
| BS25096 | 2009 | Adult | M | 2009-05-17 | 2011-03-27 | 679 | 328 |
| BS13035 | 2009 | Adult | F | 2009-05-17 | 2011-05-21 | 734 | 353 |
| BS9109 | 2009 | Adult | M | 2009-05-17 | 2011-03-07 | 659 | 212 |
| BS12380 | 2009 | Adult | M | 2009-05-17 | 2011-03-14 | 667 | 371 |
| BS13136 | 2009 | Adult | F | 2009-05-17 | 2011-02-23 | 647 | 338 |
| BS25008 | 2011 | Immature | F | 2011-03-27 | 2012-03-17 | 356 | 284 |
| BS25039 | 2011 | Immature | M | 2011-04-08 | 2015-03-06 | 1428 | 297 |
| BS22495 | 2011 | Immature | M | 2011-04-15 | 2012-02-21 | 313 | 274 |
| BS25036 | 2011 | Immature | F | 2011-04-23 | 2012-03-17 | 329 | 262 |
| BS25049 | 2011 | Immature | M | 2011-05-19 | 2012-04-25 | 342 | 212 |
| BS25004 | 2011 | Immature | M | 2011-05-25 | 2012-04-25 | 337 | 278 |
| BS25027 | 2011 | Immature | M | 2011-05-25 | 2013-06-18 | 755 | 259 |
| BS25017 | 2011 | Immature | F | 2011-05-28 | 2012-02-13 | 261 | 262 |
| BS29604 | 2011 | Juvenile | M | 2011-12-05 | 2018-03-27 | 2305 | 1090 |
| BS29605 | 2011 | Juvenile | M | 2011-12-05 | 2019-04-15 | 2689 | 1059 |
| BS29608 | 2011 | Juvenile | M | 2011-12-05 | 2015-04-20 | 1233 | 601 |
| BS29552 | 2011 | Juvenile | F | 2011-12-05 | 2018-03-16 | 2294 | 1220 |
| BS29553 | 2011 | Juvenile | F | 2011-12-05 | 2018-03-17 | 2294 | 1163 |
| BS29555 | 2011 | Juvenile | M | 2011-12-05 | 2017-06-28 | 2032 | 1125 |
| BS29556 | 2011 | Juvenile | F | 2011-12-06 | 2018-03-17 | 2294 | 1136 |
| BS29559 | 2011 | Juvenile | F | 2011-12-06 | 2020-03-13 | 3021 | 472 |
| BS29560 | 2011 | Juvenile | F | 2011-12-06 | 2017-03-01 | 1913 | 1059 |
| BS29561 | 2011 | Juvenile | F | 2011-12-06 | 2018-02-17 | 2266 | 1184 |
| BS33763 | 2012 | Immature | M | 2012-03-11 | 2013-02-15 | 342 | 340 |
| BS22456 | 2012 | Immature | M | 2012-03-11 | 2013-02-21 | 347 | 268 |
| BS25049 | 2012 | Immature | M | 2012-04-25 | 2018-03-16 | 2152 | 212 |
| BS25004 | 2012 | Immature | M | 2012-04-25 | 2013-02-15 | 296 | 278 |
| BS29502 | 2012 | Immature | NaN | 2012-05-20 | 2013-03-11 | 296 | 279 |

**Table S3.**
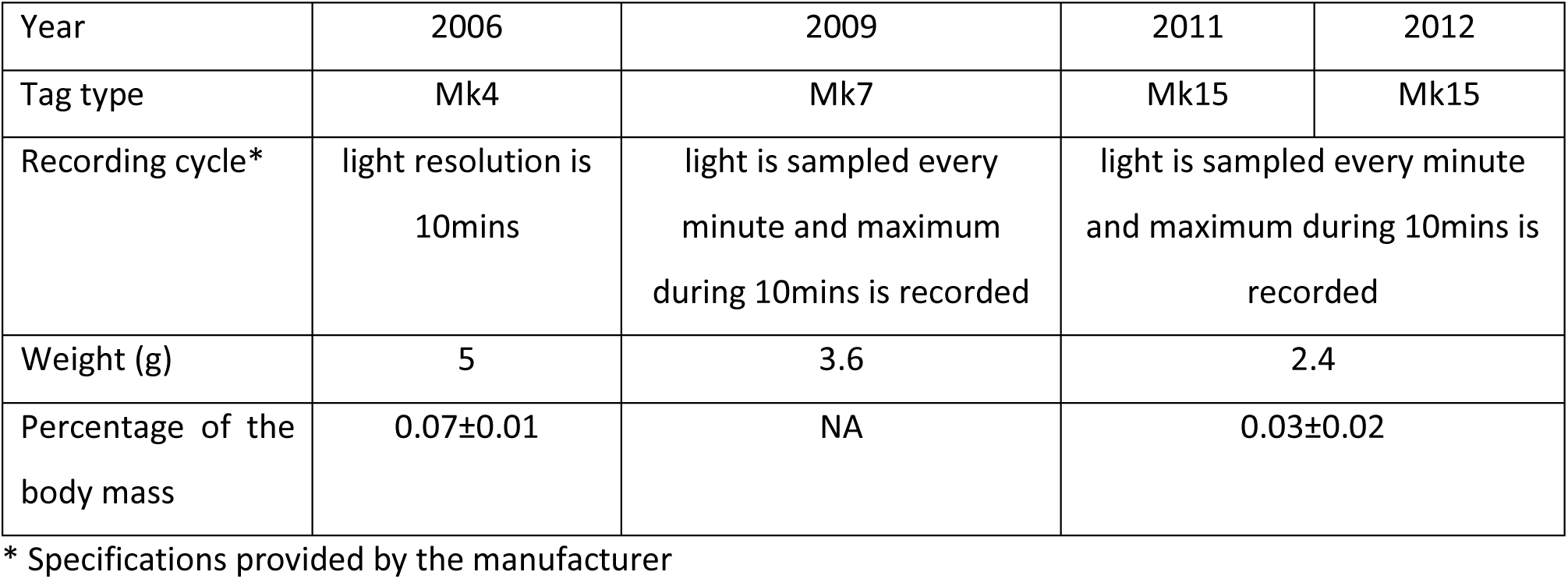
Summary of the properties of geolocators (GLS) deployed on the 36 Amsterdam albatross equipped in 2006, 2009, 2011 and 2012, at Amsterdam Island. It includes the GLS type, recording cycle, weight expressed in g, the percentage of the body mass of individuals.

| Year | 2006 | 2009 | 2011 | 2012 |
| --- | --- | --- | --- | --- |
| Tag type | Mk4 | Mk7 | Mk15 | Mk15 |
| Recording cycle* | light resolution is 10mins | light is sampled every minute and maximum during 10mins is recorded | light is sampled every minute and maximum during 10mins is recorded |  |
| Weight (g) | 5 | 3.6 | 2.4 |  |
| Percentage of the body mass | 0.07±0.01 | NA | 0.03±0.02 |  |

\* Specifications provided by the manufacturer

## REFERENCES

Adams J, Flora S (2010) Correlating seabird movements with ocean winds: linking satellite telemetry with ocean scatterometry. Marine Biology

Alerstam T, Gudmundsson GA, Larsson B (1993) Flight tracks and speeds of Antarctic and Atlantic seabirds: radar and optical measurements. Phil Trans R Soc Lond 340:55–67

Amélineau F, Péron C, Lescroël A, et al (2014) Windscape and tortuosity shape the flight costs of northern gannets. Journal of Experimental Biology 217:876–885

Arrigo KR, van Dijken GL, Bushinsky S (2008) Primary production in the Southern Ocean, 1997–2006. Journal of Geophysical Research: Oceans 113:. 10.1029/2007JC004551

Behera SK, Yamagata T (2001) Subtropical SST dipole events in the southern Indian Ocean. Geophysical Research Letters 28:327–330

Blunden J, Boyer T, Bartow-Gillies E (2023) State of the Climate in 2022. Bulletin of the American Meteorological Society 104:S1–S516. 10.1175/2023BAMSStateoftheClimate.1

Brønnvik H, Nourani E, Fiedler W, Flack A (2024) Experience reduces route selection for conspecifics by the collectively migrating white stork. Current Biology 34:2030–2037.e3. 10.1016/j.cub.2024.03.052

Cai W, Shi G, Cowan T, et al (2005) The response of the Southern Annular Mode, the East Australian Current, and the southern mid-latitude ocean circulation to global warming. Geophys Res Lett 32:L23706. 10.1029/2005GL024701

Cai W, Sullivan A, Cowan T (2009) Climate change contributes to more frequent consecutive positive Indian Ocean Dipole events. Geophysical Research Letters 36:. 10.1029/2009GL040163

Chapman JW, Klaassen RH, Drake VA, et al (2011) Animal orientation strategies for movement in flows. Current Biology 21:R861–R870

Clay TA, Joo R, Weimerskirch H, et al (2020) Sex-specific effects of wind on the flight decisions of a sexually dimorphic soaring bird. Journal of Animal Ecology 89:1811–1823

Cleveland WS, Devlin SJ (1988) Locally Weighted Regression: An Approach to Regression Analysis by Local Fitting. Journal of the American Statistical Association 83:596–610. 10.1080/01621459.1988.10478639

Clipp HL, Cohen EB, Smolinsky JA, et al (2020) Broad-scale weather patterns encountered during flight influence landbird stopover distributions. Remote Sensing 12:565

de Grissac S, Börger L, Guitteaud A, Weimerskirch H (2016a) Contrasting movement strategies among juvenile albatrosses and petrels. Scientific reports 6:26103

de Grissac S, Börger L, Guitteaud A, Weimerskirch H (2016b) Contrasting movement strategies among juvenile albatrosses and petrels. Scientific reports 6:26103

Delord K, Weimerskirch H, Barbraud C (2024) The challenges of independence: ontogeny of at-sea behaviour in a long-lived seabird. Peer Community Journal 4:. 10.24072/pcjournal.386

Dias MP, Martin R, Pearmain EJ, et al (2019) Threats to seabirds: A global assessment. Biological Conservation 237:525–537

Dokter AM, Shamoun-Baranes J, Kemp MU, et al (2013) High altitude bird migration at temperate latitudes: a synoptic perspective on wind assistance. PloS one 8:e52300

Drake V, Farrow R (1988) The influence of atmospheric structure and motions on insect migration. Annual review of entomology 33:183–210

E.U. Copernicus Marine Service Information (2012) Global Ocean Wind L4 Reprocessed 6 hourly Observations -WIND_GLO_WIND_L4_REP_OBSERVATIONS_012_006

Fisel F, Heine G, Rohde C, et al (2024) Influence of age on spatial and temporal migratory patterns of Black Storks from Germany. J Ornithol 165:861–868. 10.1007/s10336-024-02170-3

Gent PR (2016) Effects of Southern Hemisphere Wind Changes on the Meridional Overturning Circulation in Ocean Models. Annu Rev Mar Sci 8:79–94. 10.1146/annurev-marine-122414-033929

Goyal R, Sen Gupta A, Jucker M, England MH (2021) Historical and Projected Changes in the Southern Hemisphere Surface Westerlies. Geophys Res Lett 48:. 10.1029/2020GL090849

Hannachi A, Jolliffe IT, Stephenson DB (2007) Empirical orthogonal functions and related techniques in atmospheric science: A review. International Journal of Climatology 27:1119–1152. 10.1002/joc.1499

Hromádková T, Pavel V, Flousek J, Briedis M (2020) Seasonally specific responses to wind patterns and ocean productivity facilitate the longest animal migration on Earth. Marine Ecology Progress Series 638:1–12

Iacobucci A, Noullez A (2005) A Frequency Selective Filter for Short-Length Time Series. Comput Econ 25:75–102. 10.1007/s10614-005-6276-7

Jianping L, Wang JXL (2003) A new North Atlantic Oscillation index and its variability. Adv Atmos Sci 20:661–676. 10.1007/BF02915394

Johnson S (1995) Insect migration in North America: synoptic-scale transport in a highly seasonal environment. Insect migration: tracking resources through space and time Cambridge University Press, Cambridge 31–66

Jolliffe I (2011) Principal component analysis. In: International encyclopedia of statistical science. Springer, pp 1094–1096

Jouventin P, Martinez J, Roux J-P (1989) Breeding biology and current status of the Amsterdam Island Albatross Diomedea amsterdamensis. Ibis 131:182-

Kang JM, Shaw TA, Kang SM, et al (2024) Revisiting the reanalysis-model discrepancy in Southern Hemisphere winter storm track trends. npj Clim Atmos Sci 7:252. 10.1038/s41612-024-00801-3

Krishnamurti TN, Bhalme H (1976) Oscillations of a monsoon system. Part I. Observational aspects. Journal of Atmospheric Sciences 33:1937–1954

Kürten N, Wynn J, Haest B, et al (2025) Route flexibility is associated with headwind minimization in a long-distance migratory seabird. Proceedings of the Royal Society B: Biological Sciences 292:20242522. 10.1098/rspb.2024.2522

La Sorte FA, Fink D (2017) Projected changes in prevailing winds for transatlantic migratory birds under global warming. Journal of Animal Ecology 86:273–284. 10.1111/1365-2656.12624

Lionello P, D’Agostino R, Ferreira D, et al (2024) The Hadley circulation in a changing climate. Annals of the New York Academy of Sciences 1534:69–93. 10.1111/nyas.15114

Lisovski S, Hahn S (2012) GeoLight – processing and analysing light-based geolocator data in R. Methods in Ecology and Evolution 3:1055–1059. 10.1111/j.2041-210X.2012.00248.x

Lisovski S, Hewson CM, Klaassen RHG, et al (2012) Geolocation by light: accuracy and precision affected by environmental factors. Methods in Ecology and Evolution 3:603–612. 10.1111/j.2041-210X.2012.00185.x

Manola I, Bradarić M, Groenland R, et al (2020) Associations of synoptic weather conditions with nocturnal bird migration over the North Sea. Frontiers in Ecology and Evolution 8:542438

Masson-Delmotte V, Zhai P, Pirani A, et al (2021) Contribution of working group I to the sixth assessment report of the intergovernmental panel on climate change. Climate change

Morten JM, Buchanan PJ, Egevang C, et al (2023) Global warming and arctic terns: Estimating climate change impacts on the world’s longest migration. Global Change Biology

Pennycuick CJ (1982) The flight of petrels and albatrosses (Procellariiformes), observed in South Georgia and its vicinity. Phil Trans R Soc Lond 300:75–106

Pennycuick CJ (1978) Fifteen testable predictions about bird flight. Oikos 165–176

Péron C, Authier M, Barbraud C, et al (2010) Interdecadal changes in at-sea distribution and abundance of subantarctic seabirds along a latitudinal gradient in the Southern Indian Ocean. Global Change Biology 16:1895–1909

Phillips RA, Silk JRD, Croxall JP, et al (2004) Accuracy of geolocation estimates for flying seabirds. Marine Ecology Progress Series 266:265–272. 10.3354/meps266265

Powers KD, Pratte I, Ronconi RA, et al (2022) Age-Related Interactions with Wind During Migration Support the Hypothesis of Developmental Learning in a Migrating Long-Lived Seabird. Front Mar Sci 9:. 10.3389/fmars.2022.938033

Rasmusson EM, Wallace JM (1983) Meteorological Aspects of the El Niño/Southern Oscillation. Science 222:1195–1202. 10.1126/science.222.4629.1195

Richardson PL (2019) Leonardo da Vinci’s discovery of the dynamic soaring by birds in wind shear. Notes and Records: the Royal Society journal of the history of science 73:285–301

Riotte-Lambert L, Weimerskirch H (2013) Do naive juvenile seabirds forage differently from adults? Proceedings of the Royal Society B: Biological Sciences 280:20131434

Rivalan P, Barbraud C, Inchausti P, Weimerskirch H (2010) Combined impact of longline fisheries and climate on the persistence of the Amsterdam albatross. Ibis 152:6–18

Roux JP, Jouventin P, Mougin JL, et al (1983) Un nouvel albatros Diomedea amsterdamensis n. sp. découvert sur l’île Amsterdam (37°50’S, 77°35’E). L’Oiseau et R F O 53:1–11

Roy A, Désert T, Delcourt V, et al (2025) Enhanced forecasting of bird nocturnal migration intensity in relation to previous days and synoptic weather patterns. International Journal of Biometeorology

Sachs G, Traugott J, Nesterova AP, Bonadonna F (2013) Experimental verification of dynamic soaring in albatrosses. Journal of Experimental Biology 216:4222–4232

Safi K, Kranstauber B, Weinzierl R, et al (2013) Flying with the wind: scale dependency of speed and direction measurements in modelling wind support in avian flight. Movement Ecology 1, 4

Schoombie S, Wilson RP, Ryan PG (2023) Wind driven effects on the fine-scale flight behaviour of dynamic soaring wandering albatrosses. Marine Ecology Progress Series WIND: 10.3354/meps14265

Sergio F, Barbosa JM, Tanferna A, et al (2022) Compensation for wind drift during raptor migration improves with age through mortality selection. Nat Ecol Evol 6:989–997. 10.1038/s41559-022-01776-1

Shamoun-Baranes J, Liechti F, Vansteelant WM (2017) Atmospheric conditions create freeways, detours and tailbacks for migrating birds. Journal of Comparative Physiology A 203:509–529

Sharmar VD, Markina MYu, Gulev SK (2021) Global Ocean Wind-Wave Model Hindcasts Forced by Different Reanalyzes: A Comparative Assessment. Journal of Geophysical Research: Oceans 126:e2020JC016710. 10.1029/2020JC016710

Shaw TA, Arblaster JM, Birner T, et al (2024) Emerging Climate Change Signals in Atmospheric Circulation. AGU Advances 5:e2024AV001297. 10.1029/2024AV001297

Shaw TA, Miyawaki O, Donohoe A (2022) Stormier Southern Hemisphere induced by topography and ocean circulation. Proceedings of the National Academy of Sciences 119:e2123512119. 10.1073/pnas.2123512119

Skyllas N, Loonen MJJE, Bintanja R (2023) Arctic tern flyways and the changing Atlantic Ocean wind patterns. Climate Change Ecology 6:100076. 10.1016/j.ecochg.2023.100076

Somveille M, Dias MP, Weimerskirch H, Davies TE (2020) Projected migrations of southern Indian Ocean albatrosses as a response to climate change. Ecography 43:1683–1691

Spear LB, Ainley DG (1998) Morphological differences relative to ecological segregation in petrels (family:Procellariidae) of the Southern Ocean and Tropical Pacific. The Auk 115:1017–1033

Spruzen FL, Woehler EJ (2002) The influence of synoptic weather patterns on the at-sea behaviour of three species of albatross. Polar Biology 25:296–302

Swart NC, Fyfe JC (2012) Observed and simulated changes in the Southern Hemisphere surface westerly wind-stress: CHANGES IN THE S.H. WESTERLIES. Geophys Res Lett 39:n/a-n/a. 10.1029/2012GL052810

Sydeman W, García-Reyes M, Schoeman DS, et al (2014) Climate change and wind intensification in coastal upwelling ecosystems. Science 345:77–80

Terray P (2011) Southern Hemisphere extra-tropical forcing: a new paradigm for El Niño-Southern Oscillation. Climate dynamics 36:2171–2199

Thiebot J-B, Delord K, Barbraud C, et al (2016) 167 individuals versus millions of hooks: bycatch mitigation in longline fisheries underlies conservation of Amsterdam albatrosses. Aquatic Conservation: Marine and Freshwater Ecosystems 26:674–688. 10.1002/aqc.2578

Thiebot J-B, Delord K, Marteau C, Weimerskirch H (2014) Stage-dependent distribution of the critically endangered Amsterdam albatross in relation to Economic Exclusive Zones. Endangered Species Research 23:263–276

Thorne LH, Clay TA, Phillips RA, et al (2023) Effects of wind on the movement, behavior, energetics, and life history of seabirds. Marine Ecology Progress Series 723:73–117. 10.3354/meps14417

Thorne LH, Conners MG, Hazen EL, et al (2016) Effects of El Niño-driven changes in wind patterns on North Pacific albatrosses. J R Soc Interface 13:20160196. 10.1098/rsif.2016.0196

Van Doren BM, Horton KG (2018) A continental system for forecasting bird migration. Science 361:1115–1118

Vansteelant WMG, Gangoso L, Bouten W, et al (2021) Adaptive drift and barrier-avoidance by a fly-forage migrant along a climate-driven flyway. Movement Ecology 9:37. 10.1186/s40462-021-00272-8

Ventura F, Granadeiro JP, Padget O, Catry P (2020) Gadfly petrels use knowledge of the windscape, not memorized foraging patches, to optimize foraging trips on ocean-wide scales. Proceedings of the Royal Society B-Biological Sciences 287:20191775. 10.1098/rspb.2019.1775

Weimerskirch H, Louzao M, de Grissac S, Delord K (2012) Changes in Wind Pattern Alter Albatross Distribution and Life-History Traits. Science 335:211–214

Weimerskirch H, T. G, J. M, et al (2000) Fast and fuel efficient? Optimal use of wind by flying albatrosses. Proceedings of the Royal Society B267:1869–1874

Wheeler MC, Hendon HH (2004) An All-Season Real-Time Multivariate MJO Index: Development of an Index for Monitoring and Prediction. Monthly Weather Review 132:1917–1932. 10.1175/1520-0493(2004)132%253C1917:AARMMI%253E2.0.CO;2

Young IR, Ribal A (2019) Multiplatform evaluation of global trends in wind speed and wave height. Science 364:548–552. 10.1126/science.aav9527

Zhao Y, Wen Z, Li X, et al (2023) Meridional variation of the Mascarene High and atmospheric transient eddy dynamical forcing over the Southern Indian Ocean in austral winter. Journal of Climate 36:6937–6950

## REFERENCES

de Grissac S, Börger L, Guitteaud A, Weimerskirch H (2016) Contrasting movement strategies among juvenile albatrosses and petrels. Scientific reports 6:26103

Delord K, Barbraud C, Bost CA, et al (2013) Atlas of top predators from the French Southern Territories in the Southern Indian Ocean. http://www.cebc.cnrs.fr/ecomm/Fr_ecomm/ecomm_ecor_OI1.html

Nevitt GA, Losekoot M, Weimerskirch H (2008) Evidence for olfactory search in wandering albatross, Diomedea exulans. Proceedings of the National Academy of Sciences 105:4576–4581

Pajot A, Corbeau A, Jambon A, Weimerskirch H (2021) Diel at-sea activity of two species of great albatrosses: the ontogeny of foraging and movement behaviour. Journal of Avian Biology 52:

Weimerskirch H, Brothers N, ouventin P (1997) Population dynamics of Wandering Albatross *Diomedea exulans* and Amsterdam Albatross *D. amsterdamensis* in the Indian Ocean and their relationship with long-line fisheries: conservation implications. Biological conservation 79:257–270

Weimerskirch H, Delord K, Guitteaud A, et al (2015) Extreme variation in migration strategies between and within wandering albatross populations during their sabbatical year, and their fitness consequences. Scientific Reports 5:8853-

Wilson RP, Ducamp JJ, Rees G, et al (1992) Estimation of location: global coverage using light intensity. In: Priede IMSS (ed). Ellis Horward, Chichester, pp 131–134

